# CX3CR1 modulates migration of resident microglia towards brain injury

**DOI:** 10.1101/2024.09.23.614458

**Authors:** Jens Wagner, Cornelia Hoyer, Henrike Antony, Kristiina Lundgrén, Rabah Soliymani, Sophie Crux, Lena Justus, Kevin Keppler, Julia Steffen, Christian Kurts, Daniel R. Engel, Jochen Herms, Maciej Łałowski, Martin Fuhrmann

## Abstract

Microglia are innate immune cells of the central nervous system (CNS). They extend their processes towards and migrate towards injuries *in vivo*. However, whether the fractalkine receptor (CX3CR1) influences microglial migration remains unknown. Label-free proteomic profiling predicted changes in RHO-signaling activity that hint at dysregulated cytoskeleton signaling in *Cx3cr1-*deficient murine cortex tissue. To further investigate microglial migration, we carried out 4-hour interval two-photon *in vivo* imaging for 72 hours after a laser lesion in the cortex. *Cx3cr1*-deficient microglia showed enhanced migration towards the lesion. Additionally, length and velocity of microglial fine processes extending towards the lesion were increased in *Cx3cr1-*deficient microglia. Migration remained unchanged in *Ccr2*-deficient mice, indicating that monocyte-derived macrophages/microglia did not contribute to microglia accumulation around the lesion. These results demonstrate microglia migration towards CNS injury and suggest CX3CR1 as a modulator of this. Manipulating microglia migration via CX3CR1 therefore is a potential target for treatment of CNS-injury.

## Introduction

Microglia are mononuclear phagocytes that reside in the parenchyma of the central nervous system^1^. During embryogenesis, yolk sac-derived microglia colonize the brain, while after this critical period microglia persist in the CNS without substantial input from bone-marrow derived myeloid cell populations under healthy conditions anymore^2–4^. It is hypothesized that resident microglia proliferate to maintain a stable population. However, under specific neurodegenerative conditions myeloid precursors may enter the brain via an impaired blood brain barrier (BBB)^5^. Microglia are highly motile cells with long processes that constantly scan the surrounding brain parenchyma^6^.

Microglia respond to injuries in the CNS. In the healthy brain, only 5% of the microglia are mobile, however, this number increases in a sex-dependent manner as a response towards brain bleeding^7^. After brain injury, microglia extend their processes towards the lesion. Within the first 30 minutes after a laser lesion, processes of microglial cells in close proximity reach the damaged site and within 1–3-hours most of them are directed towards the damage, while the microglial soma remains at its position for at least 10 hours. This migration process is driven by adenosine triphosphate (ATP) via the purinergic G protein-coupled P2Y receptor^8^. The metabotropic P2Y12 receptor is suggested as the major site where ATP induces microglial chemotaxis^9^. Indeed, ATP which is released at the site of brain damage, induces microglial migration^10,11^. Further, microglial accumulation in and around a diphtheria toxin A-chain- expression-induced lesion was shown^12^. Moreover, neuronal lesions in the brain do not induce migration of peripheral cells into the brain, concluding that the BBB remains uncompromised. Therefore, the cells accumulating in and around the lesion are microglia and not peripheral myeloid cells, such as monocytes or macrophages^12^. Not only at sites of brain injury, accumulation of microglia is detected, but also around lesions of Multiple Sclerosis (MS) pathology^13,14^. In MS, microglia are also associated with a higher lesion load^15^.

One key receptor that is involved in microglial migration and chemotaxis, is the fractalkine receptor (CX3CR1) and its ligand fractalkine (CX3CL1). Specifically, LPS-stimulated intracranially injected *Cx3cr1^−/−^* microglia do not migrate along white matter tracts^16^. In addition, increased microglia migration towards dying neurons in an AD mouse model is rescued by *Cx3cr1* knockout, further supporting a role for CX3CR1 in migration^17^. Thus, microglial *Cx3cr1*- deficiency can be either detrimental or beneficial depending on the lesion context^16–18^. Directed migration of Cx3cr1-eGFP labeled microglia towards a microhemorrhage over a time-interval of 48h *in vivo* has been previously shown^19^, however, the role of *Cx3cr1* in regulating microglial movements remains elusive.

To better understand the role of *Cx3cr1* in microglial dynamics upon brain injury under sterile inflammatory conditions, we investigated *Cx3cr1*-mediated microglial migration and proliferation upon a laser lesion using two-photon *in vivo* imaging. We performed unprecedented temporal resolution 4-hour time-interval imaging for a period of up to 72 hours to monitor microglial dynamics. We also analyzed, whether peripheral monocyte derived macrophages and proliferating microglia contributed to local accumulation of cells around the lesion. Our results support the conclusion that primarily resident microglia accumulate at a lesion, and CX3CR1 modulates microglial migration and process motility.

## Results

### Proteomics reveal differentially expressed proteins in cytoskeleton signaling

To identify potential protein expression changes in relation to CX3CR1, we carried out proteomics comparing *Cx3cr1^−/−^* and *Cx3cr1^+/–^*mice. Therefore, we performed label-free quantitative global proteomic profiling to elucidate the alterations in protein abundance between the two groups, as well as to indicate pathways/local networks that were altered based on the differential protein expression. Whole-cortex tissue of *Cx3cr1^−/−^*and *Cx3cr1^+/–^* mice was processed and proteins with differential abundance (DEPs) were determined via tandem mass spectrometry **(Fig. 1a, Supplementary Table 1)**. We performed principal component analysis (PCA) on the total count of detected proteins from each sample (three technical replicates per animal) to elucidate the degree of similarities between the groups. The PCA revealed that the most animals clustered together, but some of them were better separated leading to two overlapping clusters between the two genotypes (**Fig. 1b**). Interestingly, the proteomes of animals from the two genotypes differed significantly between the experimental groups based on analysis using an unpaired t-test as described in the methods section, hinting at functional differences (**Fig. 1c, Supplementary Table 1**). Based on the DEPs shown in the volcano plots (**Fig. 1c**), we performed Ingenuity Pathway Analysis (IPA) to predict alterations in cellular signaling processes (**Supplementary Table 2**). Out of all detected 170 canonical pathways with allotted z-score, 10 pathways related to changes in cellular migration/cytoskeleton alterations were selected (**Fig. 1d**). Indeed, we observed significant changes in Ras homology family (RHO)-signaling pathways involving cytoskeletal changes between the groups indicating that many migration-regulating signaling processes were downregulated in *Cx3cr1^−/−^* compared to *Cx3cr1^+/–^* mice. In parallel, an inhibitory regulator of RHO-signaling (RHGODI) was upregulated. RHO-specific guanine nucleotide dissociation inhibitors (RHOGDIs) maintain the inactive fraction of the RHO-GTPases in the cytosol, while only a few are active in the cytoplasm^20^. RHO-GTPases were found to modulate many cellular processes including migration as well as cytoskeleton regulation^21^. Therefore, an upregulation of RHOGDI signaling in association with downregulation of RHO-signaling and cytoskeleton modulation is a consistent finding.

**Figure 1.**
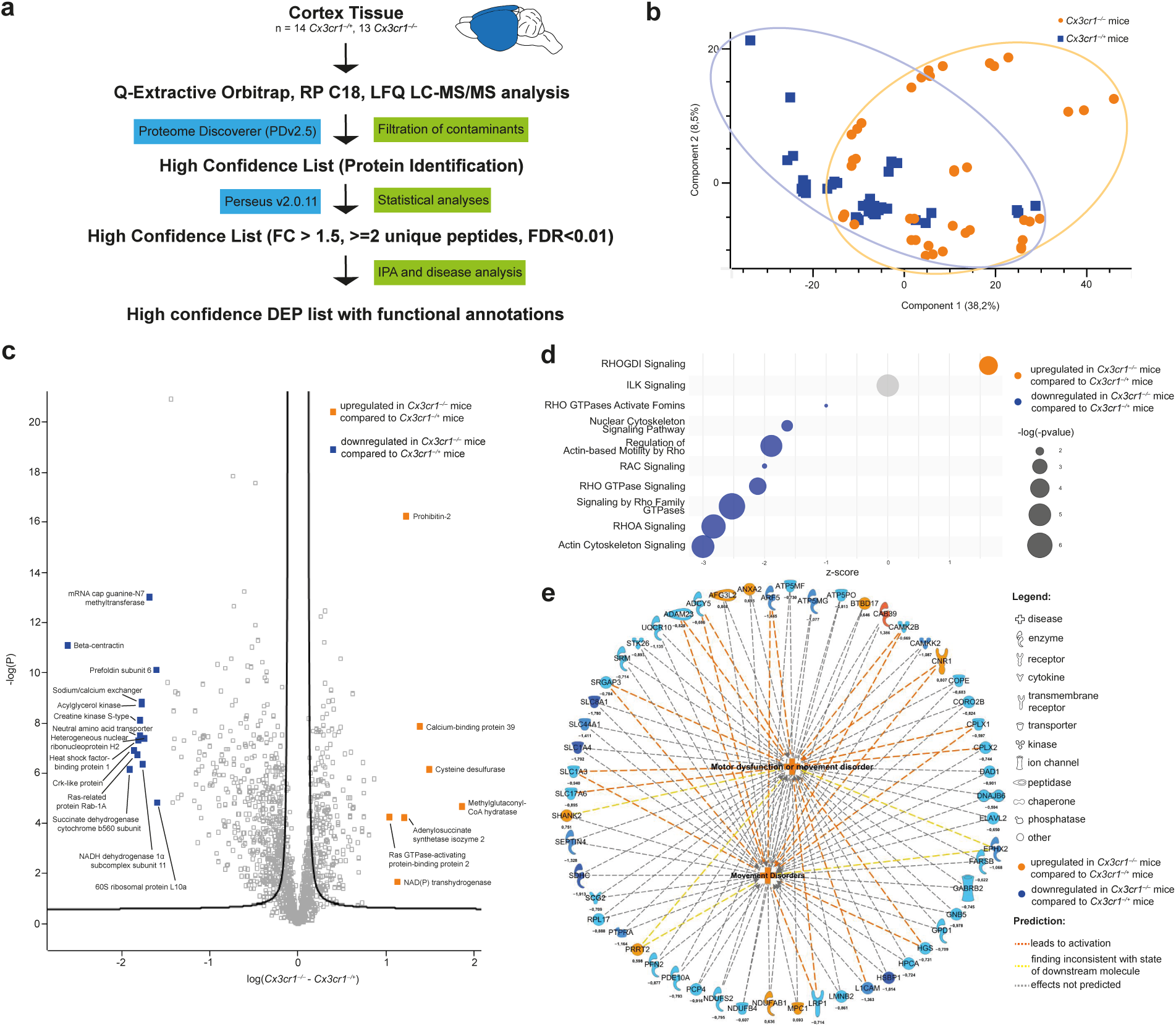
Proteomic analysis comparing *Cx3cr1^+/–^* to *Cx3cr1^−/−^* mice. **(a)** Workflow of LFQ proteomics analysis to determine alterations in protein abundance between *Cx3cr1^−/−^* and *Cx3cr1^+/–^* mice. The number of used animals (n) is indicated for each genotype. **(b)** Unsupervised PCA was performed to compare the overall separation between *Cx3cr1^−/−^* (orange) and *Cx3cr1^+/–^* (blue) mouse groups. The samples were run in technical replicates and therefore are resulting in three points per animal. **(c)** Volcano plot of the DEPs comparing *Cx3cr1^−/−^*and *Cx3cr1^+/–^* mice. The log-transformed values are plotted against the negative log of the p-value to display the statistical differences. **(d)** The chart displays all pathways related to RHO-signaling/cell-migration as determined by IPA software. The pathway alterations are predicted based on DEPs. The bubble size is according to the statistical significance of the pathway with a given -log(-p) value, and the color represents the experimental condition. The z-score value relates to activation or inhibition of a given pathway.**(e)** Predicted interactions are depicted in the network. The links between the *Cx3cr1* genotype and motor dysfunction is displayed. The arrows show the directionality of the changes. The numbers below the DEPs refer to the respective fold-changes in log scale. n = 42 14 *Cx3cr1^− /+^* and 39 *Cx3cr1^−/−^* replicas from N = 14 *Cx3cr1^−/+^* and 13 *Cx3cr1^−/−^* mice.

Since CX3CR1 is expressed only on microglia in the cortex, it implies a specific modulation of microglial cytoskeletal signaling, potentially changing migratory properties. Interestingly, previous reports have found that ablation of Rho-signaling by a RhoA-GTPase-KO in microglia triggers spontaneous microglial activation^22^.

In addition to the canonical pathway analysis, we also analyzed the interplay of the differentially regulated proteins and the predicted outcome for the organism (**Fig. 1e**). The disease/function network analysis is based on the entire proteomic profile as revealed in the analysis and showed that dysregulated DEPs in *Cx3cr1^−/−^* mice were strongly connected with predicted elevation of motor dysfunction and movement disorders. As we found a significant down- regulation in Rho-signaling, likely related to microglia, we assumed a change in microglial functionality likely resulting in changes in their mobility. Thus, we conducted further functional analysis in the murine cortex to assess how specifically microglia migration is affected by the different *Cx3cr1*-genotypes.

### CX3CR1 modulates migration and process extension of microglia

To investigate the migration of microglia around a laser lesion in the brain *in vivo*, we carried out repetitive two-photon *in vivo* imaging of individual GFP-expressing microglia over a period of 72 hours with 4–6-hour time intervals **(Fig. 2a, Supplementary Video 1)**. *Cx3cr1^gfp^* mice were used. Green fluorescent protein (GFP) is inserted into the chemokine receptor *Cx3cr1*- locus leading to *Cx3cr1*-deficiency in homozygous *Cx3cr1^gfp/gfp^*mice. As controls, heterozygous *Cx3cr1^gfp/+^* mice were used as they still feature labelled microglia while maintaining a portion of CX3CR1. Microglial GFP-expression labels the microglia for *in vivo* microscopy and imaging of the microglia movements **(Fig. 2b, c and d, Supplementary** Fig. 2).

**Figure 2.**
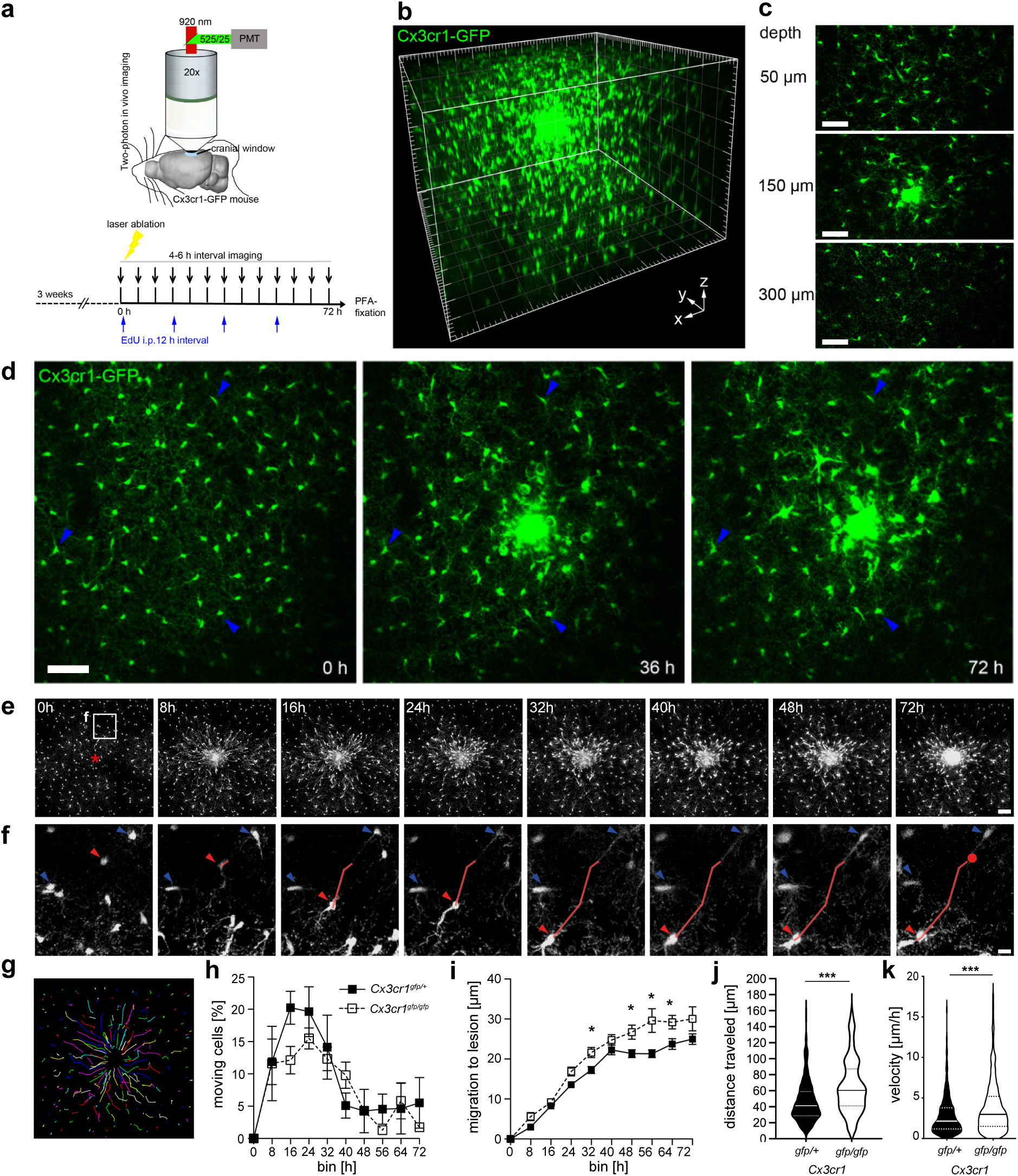
Microglia migration towards a laser lesion is modulated by CX3CR1. **(a)** Schematic of a mouse fixed under a two-photon laser scanning microscope and the timeline of the experiments. The mice used for imaging of microglia migration underwent cranial window surgery over the right somatosensory cortex to install a circular coverslip (0 5 mm) as described. **(b)** 3-dimensional reconstruction of a 300 µm z-stack showing microglia (green) around the lesion. Major tick = 50 µm **(c)** Optical slices from various depths (50 - 300 µm) illustrating microglia accumulation around the lesion. Scale bars = 50 µm **(d)** Maximum intensity projection of a laser lesion illustrating three time-points before and after the lesion. Blue arrows indicate non-moving microglia. Scale bar: 50 µm. **(e)** Time-lapse two-photon *in vivo* images of cortical microglia before and up to 72 hours after laser lesion induction. The red star marks the position of the lesion. Scale bar = 75 µm **(f)** Zoom-in from white inset in (e) illustrating a migrating microglial cell (red arrow) and non-moving microglia (blue arrows) over the 72 hours imaging period. The cell movement is traced over the imaging period (red line) with the red dot representing the starting point. Scale bar = 15 µm **(g)** Color-coded tracks of migrating and sessile microglia in (e). **(h)** The fraction of migrating microglia over time of *Cx3cr1^gfp/+^* versus *Cx3cr1^gfp/gfp^* is not different between the two genotypes (p > 0.05; determined using Two-way ANOVA followed by Šídák’s multiple comparisons test). **(i)** Distance traveled towards the lesion over time is significantly increased in *Cx3cr1^gfp/gfp^* compared to *Cx3cr1^gfp/^*^+^ mice (* p < 0.05; determined using Two-way ANOVA followed by Šídák’s multiple comparisons test). **(j)** Overall distance of microglia traveled towards the lesion is significantly increased in *Cx3cr1^gfp/gfp^* mice. (n = 351 cells in *Cx3cr1^gfp/gfp^* and 159 cells in *Cx3cr1^gfp/+^* mice *Cx3cr1^gfp/+^*: 46.19 ±1.345 µm/h; *Cx3cr1^gfp/gfp^*: 65.20 ±2.691 µm/h; *** p < 0.001; determined using an unpaired t-test) **(k)** Microglia migration velocity is significantly increased in *Cx3cr1^gfp/gfp^*in comparison to *Cx3cr1^gfp/+^* mice (n = 719 cells in *Cx3cr1^gfp/gfp^*and 461 cells in *Cx3cr1^gfp/+^* mice *Cx3cr1^gfp/+^*: 3.1 ±0.1 µm/h; *Cx3cr1^gfp/gfp^*: 4.3 ±0.2 µm/h; *** p < 0.001; determined using an unpaired t-test). (h-k) N = 5 *Cx3cr1^gfp/gfp^* and 7 *Cx3cr1^gfp/+^* mice.

Initially, we investigated microglial cell body migration starting 3-4 hours after laser lesion induction **(Fig. 2e and f)**. We tracked on average 166 ±10 microglia per mouse and lesion **(Fig. 2g)**. The majority of cells (74.0 ±3.5%) remained at their initial position in the brain parenchyma, while a substantial proportion of 26.0 ±3.5% started to migrate. The fraction of migrating cells, however, was not constant over time, it increased rapidly towards a peak (at 21 ±3h after the laser lesion and declined to 4.3 ±0.8% moving cells after 48 hours **(Fig. 2h)**. Time to peak was not different between *Cx3cr1^gfp/+^*and *Cx3cr1^gfp/gfp^* mice **(Fig. 2h)**. Surprisingly, the distance traveled towards the lesion was significantly increased in *Cx3cr1^gfp/+^* compared to *Cx3cr1^gfp/gfp^* microglia **(Fig. 2i and j)**. Additionally, *Cx3cr1^gfp/gfp^* microglia exhibited a significant 41% increase in migration velocity in comparison to *Cx3cr1^gfp/+^* microglia **(Fig. 2k)**. The lesion diameter was the same in both experimental groups, ruling out any biasing effect due to different lesion size in chemotactic gradient **(Supplementary** Fig. 1a). Simultaneously, recruitment radius and microglial density remained unchanged between the two conditions **(Supplementary** Fig. 1b, d and e). Moreover, MI and total percentage of migrating microglia was unaffected by genotype, indicating that *Cx3cr1^gfp/gfp^* microglia were able to migrate, but they migrated further with higher speed **(Supplementary** Fig. 1c and f). Interestingly, also microglia motility, the turnover rate of the microglia fine processes, remains unaffected between the conditions indicating that surveillance capacity as well as not migration-related functions are unaffected by *Cx3cr1* **(Supplementary** Fig. 3). Taken together, migration distance and velocity of fine processes are enhanced in *Cx3cr1^gfp/gfp^* microglia, indicating a crucial role of CX3CR1 in regulating dynamics in response to brain injury.

In addition to microglial cell body migration, we analyzed the effect of *Cx3cr1*-deficiency on the kinetics of microglia processes’ extension towards a laser lesion. We repetitively imaged in 5- minute time intervals throughout the first hour after lesion induction **(Fig. 3a and b; Supplementary Video 2)**. As previously shown, microglial fine processes extended towards the lesion earlier than the cell body **(Fig. 2e, 3a and b)** indicating that processes extension precedes cell body migration.

**Figure 3.**
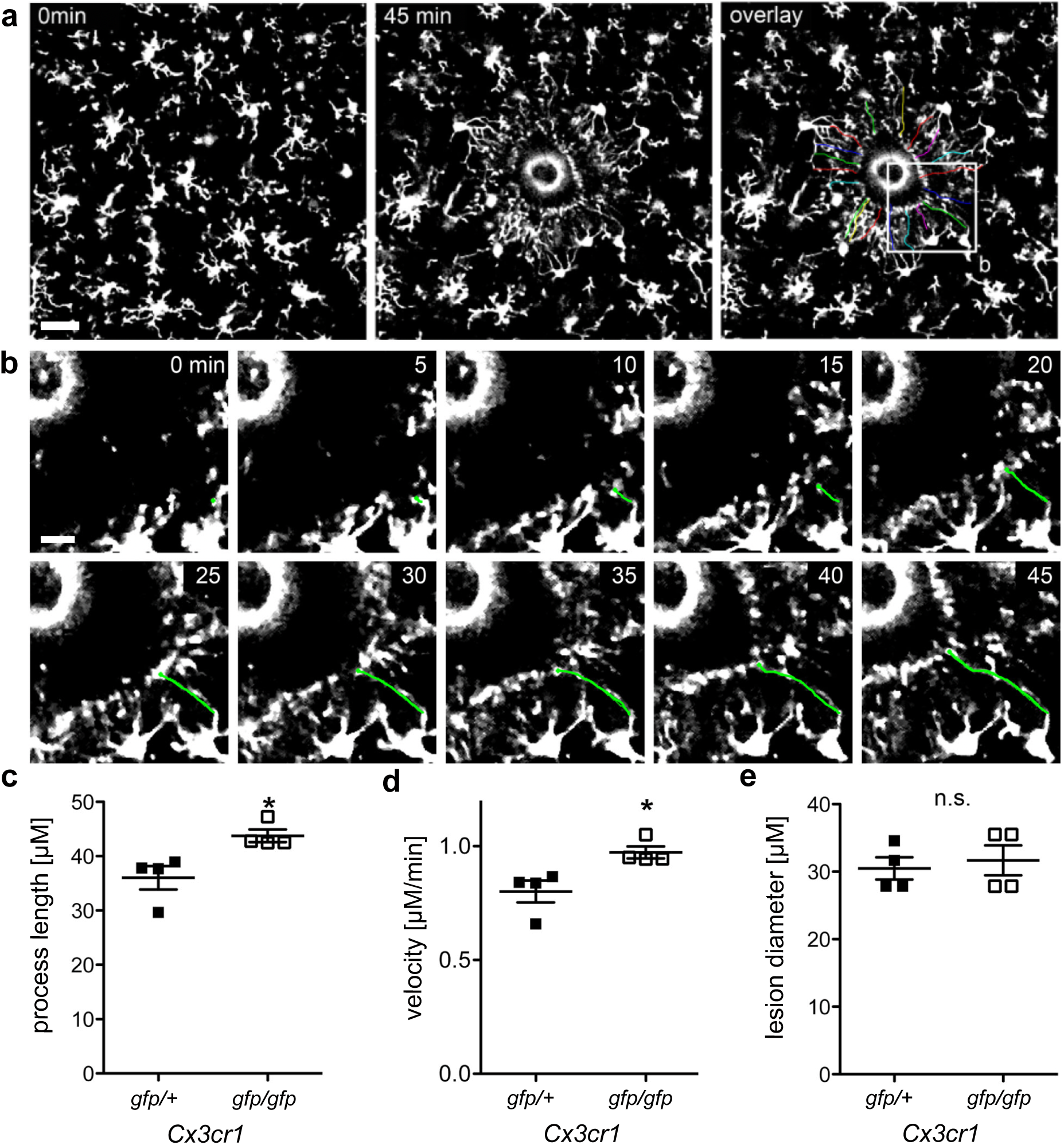
Microglial process extension is modulated by CX3CR1. **(a)** Exemplary overview z-plane of microglia before (0 min) and after a laser lesion (45 min). The third panel illustrates individual, tracked fine processes (color-coded lines). Scale bar = 25 µm **(b)** Enlarged image of the highlighted section in (a) showing an individual microglial fine process marked by a green track from 0-45 min in 5 min intervals. Scale bar = 10 µm **(c, d)** The fine processes length (c) and average velocity (d) of microglia fine processes extending towards the lesion comparing *Cx3cr1^gfp/gfp^* mice (fine process length: *Cx3cr1^gfp/+^*: 36.0 ±2.1 µm; *Cx3cr1^gfp/gfp^*: 43.8 ±1.2 µm; n = 4 mice/group; * p < 0.05; velocity: *Cx3cr1^gfp/+^*: 0.80 ±0.05 µm/min; *Cx3cr1^gfp/gfp^*: 0.97 ±0.03 µm/min). **(e)** Comparison of the lesion diameter between *Cx3cr1^gfp/+^* and *Cx3cr1^gfp/gfp^* mice (lesion diameter: *Cx3cr^gfp/+^*: 31 ±2 µm; *Cx3cr1^gfp/gfp^*: 32 ±2 µm; n = 4 mice/group; p > 0.05). Data in (c, d and e) are displayed as individual measurements, depicting the mean ± SEM.

Similarly to cell body migration, *Cx3cr1^gfp/gfp^* microglia have significantly longer processes, extending towards the lesion, than *Cx3cr1^gfp/+^*microglia **(Fig. 3c)**. Similarly, the velocity of process extension is significantly augmented **(Fig. 3d)**, whereas the average diameter of the laser lesion was not different between the two genotypes **(Fig. 3e)**. In summary, fine processes extension resembles the same pattern as previously described for microglia cell body migration. These data underscore the importance of CX3CR1 in facilitating microglial dynamics.

### Resident microglia migrate to the lesion and proliferate

We observed an increase of microglia density in close proximity to the lesion (r < 140 µm), a decrease within a radius of 140 to 250 µm, and a constant density more than 250 µm away from the lesion **(Fig. 4a and b)**. To decipher the origin of the microglia in close proximity to the lesion, we followed individual microglia over-time up to 72 hours with 4–6-hour acquisition intervals **(Supplementary Video 1)**. Strikingly, our observations revealed that the majority (∼98%) of microglia accumulating around the lesion were previously sessile in the brain parenchyma and started to migrate towards the lesion **(Fig. 2h)**. We occasionally observed newly appearing cells in close proximity to a neighboring microglial cell **(Fig. 4c; Supplementary Video 3)**. In order to identify if these microglia originated from cell division, we injected the proliferation marker 5-ethynyl-2’-deoxyuridine (EdU) during the 72-hour *in vivo* imaging period **(Fig. 2a)**. After imaging, we retrieved putative dividing microglia in PFA-fixed brain slices and demonstrated the division by EdU staining. As visualized, cell division was in some cases preceded by an increase in microglial soma volume **(Fig. 4c)**. We analyzed the spatio-temporal distribution of dividing microglia dependent on *Cx3cr1*-deficiency. In comparison to the fraction of migrating microglia (26.0 ±3.5%) we only detected about 1.8% dividing microglia that contributed to local cell accumulation around the laser lesion **(Fig. 4d)**. Divisions took place 49 ±2h in *Cx3cr1^gfp/+^* compared to 40 ±4h in *Cx3cr1^gfp/gfp^* mice at a time when the peak of migration had already been reached **(Fig. 4e)**. In addition, division close to the lesion is independent of CX3CR1 **(Fig. 4f)**. Interestingly, cell proliferation started after microglia processes extension and migration to the lesion **(Fig. 4e)** and therefore is not the first response towards brain parenchyma injury. Concluding, the observed migrating microglia are initially static and resident in the brain parenchyma. They start at moving about 8 hours after lesion induction. After migration towards the lesion, a small fraction of microglia proliferates in close proximity to the lesion. Local proliferation is independent of CX3CR1.

**Figure 4.**
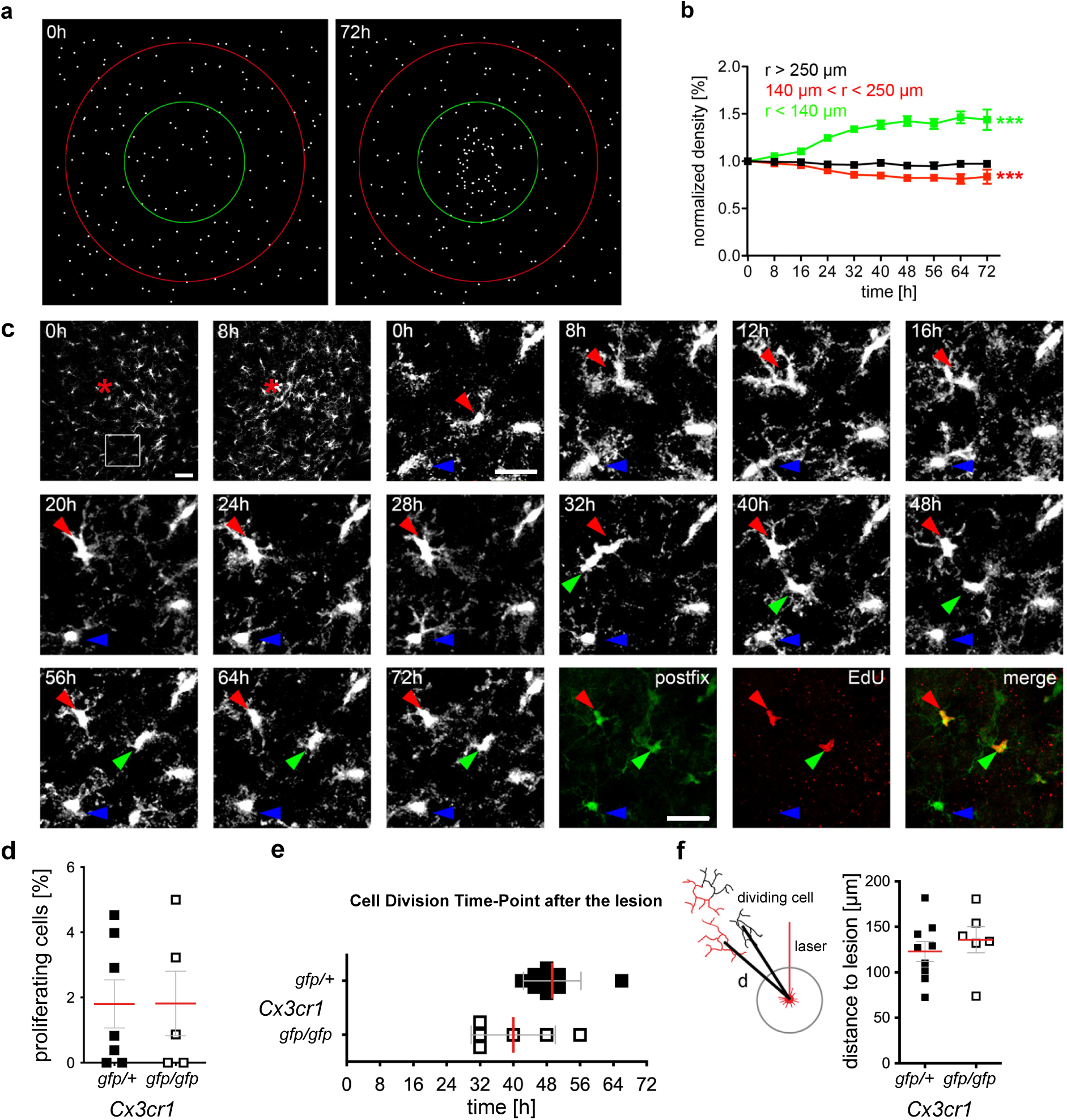
Microglia migration and proliferation of resident microglial populations. **(a)** Recording of microglia density changes around the lesion. The microglia distribution (each dot represents a tracked microglia) in the same mouse at the same cortical position before (0h) and at 72 hours (72h) after a laser lesion is shown. Red circle: radius of 250 µm, green circle: radius of 140 µm. Red star: position of the lesion **(b)** Microglia density in percent normalized to the laser lesion (r = 0). The density of microglia around the lesion (r < 140 µm) increases significantly, while the density in the region between 140-250 µm radius decreases significantly. The microglia density beyond (r > 250 µm) stays stable over time (n = 11 mice; *** p < 0.001; determined using Two-Way ANOVA). **(c)** Overview (first two images, time-point 0 and 8h) and zoom-in after perfusion and relocation (time-point 72h) of a resident dividing microglial cell (red, green arrow) after a laser lesion. The blue arrow indicates a non-dividing microglial cell. After *in vivo* imaging, microglia were re-localized in PFA-fixed brain slices and stained for the proliferation marker EdU (last three images). Scale bars: 50 µm (overviews), 25 µm (zoom-ins). **(d)** The fraction of proliferating microglia averaged over mice was independent of *Cx3cr1*-deficiency (*Cx3cr1^gfp/+^*: 1.8 ±0.7 %; *Cx3cr1^gfp/gfp^*: 1.8 ±1.0 %; p > 0.05; determined using an unpaired t-test). **(e)** Microglia proliferation events took place about 40 hours after induction of the laser lesion and independent of *Cx3cr1* (*Cx3cr1^gfp/+^*: 49 ±2 h; *Cx3cr1^gfpgfp^*: 40 ±4 h; p > 0.05; determined using an unpaired t-test). **(f)** Prior to division, microglia migrate towards the lesion (red line indicates the laser) and proliferate in proximity to the lesion (distance d) unaffected by *Cx3cr1* knockout (*Cx3cr1^gfp/+^*: 123 ±11 µm; *Cx3cr1^gfp/gfp^*: 136 ±14 µm; p > 0.05; determined using an unpaired t-test). N = 5 *Cx3cr1^gfp/gfp^*and 7 *Cx3cr1^gfp/+^* mice.

### Accumulating microglia do not originate from infiltrating monocytes

Transient recruitment of *Cx3cr1*-expressing inflammatory monocytes from the blood has been shown to depend on the chemokine receptor 2 (CCR2) expression in a model of experimental autoimmune encephalitis (EAE)^23^. And, monocyte-derived cells have been found to enter the brain under certain circumstances^24^. Our result of resident microglia migrating and accumulating at the lesion site indicates that contribution of blood-derived monocytes might be limited. To rule out that *Cx3cr1*-positive monocytes were contributing to the accumulating cells at the lesion, we used *Ccr2*-KO mice, where monocyte recruitment from blood is prevented. Therefore, we crossbred *Ccr2*-deficient mice with *Cx3cr1^gfp^* mice to yield *Cx3cr1^gfp/+^Ccr2*^−/−^ mice and carried out a laser lesion experiment as described before. The fraction of migrating microglia in *Cx3cr1^gfp/+^Ccr2^+/+^* mice changes in a similar manner over the imaging time-period with a peak of microglia migration at 16h post lesion induction **(Fig. 5b)**. We did not detect any significant differences in the proportion of migrating cells comparing *Cx3cr1^gfp/+^Ccr2^+/+^*and *Cx3cr1^gfp/+^Ccr2*^−/−^ **(Fig. 5a)**. Moreover, the kinetics, the distance traveled, the recruitment radius and the velocity remain unchanged between the conditions **(Fig. 5c, d and e)**. These results suggest that infiltrating monocyte precursors of macrophages do not seem to contribute to the accumulating of cells around the laser lesion. In addition, these data indicate that microglial chemotaxis towards the laser lesion is independent of the monocyte chemoattractant protein-1 (MCP1), the ligand for CCR2.

**Figure 5.**
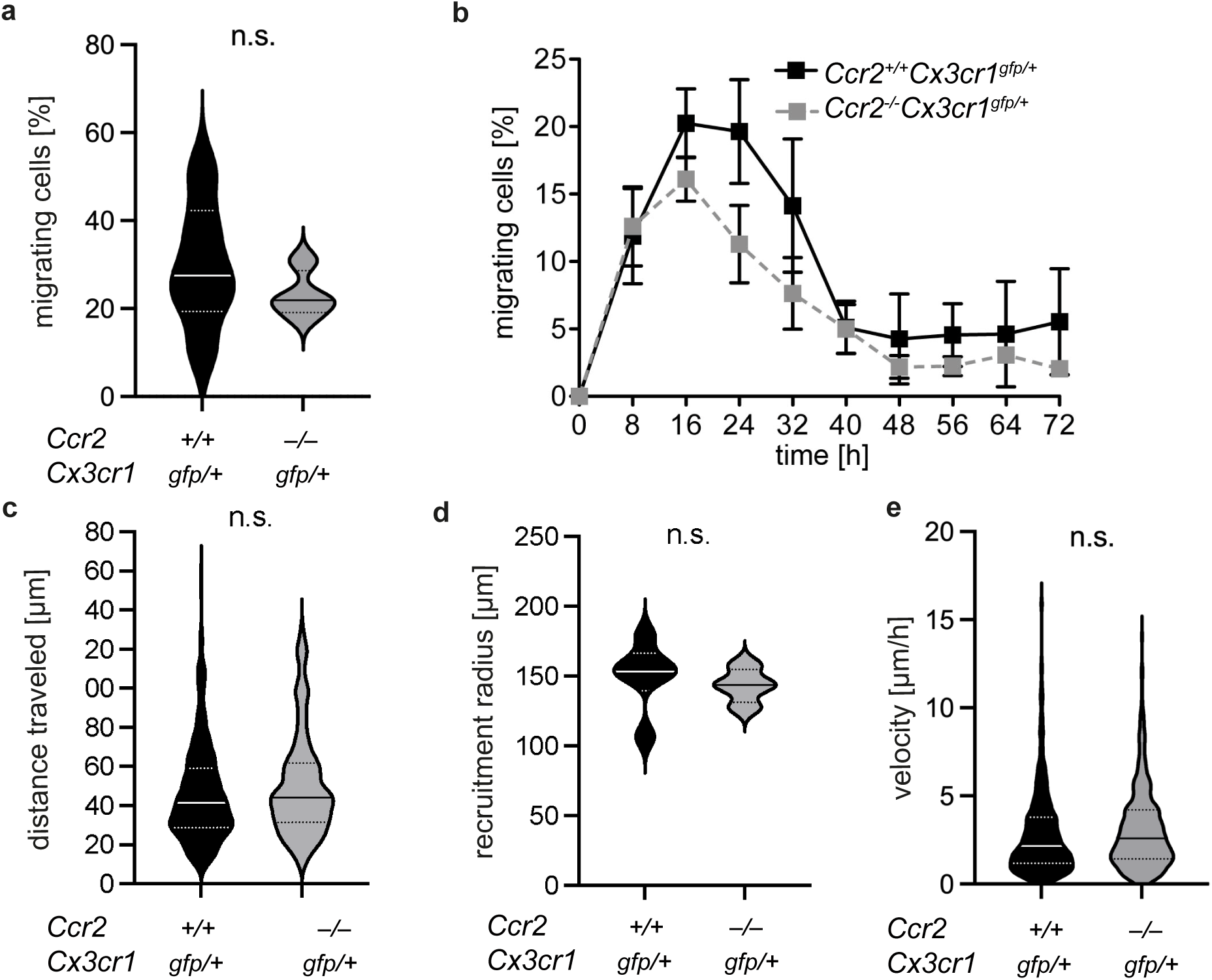
Microglia migration is independent of CCR2. **(a, b)** The fraction of migrating microglia cells overall (a) and over time (b) was not different between *Ccr2^+/+^Cx3cr1^gfp/+^* and *Ccr2^−/−^Cx3cr1^gfp/+^*mice (*Ccr2^+/+^Cx3cr1^gfp/+^*: 29.8 ±5.8% versus *Ccr2^−/−^Cx3cr1^gfp/+^*: 23.2 ±2.7%; n= 4-6 mice/group; p > 0.05; Two-way ANOVA followed by Šídák’s multiple comparisons test). **(c)** The distance traveled of microglia remained unchanged between *Ccr2^+/+^Cx3cr1^gfp/+^* and *Ccr2^−/−^Cx3cr1^gfp/+^* mice (*Ccr2^+/+^Cx3cr1^gfp/+^*: 42.7 ±2.0 µm*; Ccr2^− /−^Cx3cr1^gfp/+^*: 43.9 ±2.1 µm; p > 0.05; n = 167 cells from *Ccr2^−/−^Cx3cr1^gfp/+^* and 351 cells from *Ccr2^+/+^Cx3cr1^gfp/+^*mice; determined using an unpaired t-test). **(d)** The average recruitment radius was independent of *Ccr2*-deficiency (*Ccr2^+/+^Cx3cr1^gfp/+^*: 151 ±10 µm; *Ccr2^−/−^Cx3cr1^gfp/+^*: 143 ±6 µm; p > 0.05; determined using an unpaired t-test). **(e)** Microglia velocity was independent of *Ccr2* knockout (*Ccr2^+/+^Cx3cr1^gfp/+^*: 3.2 ±0.1 µm/h; *Ccr2^−/−^Cx3cr1^gfp/+^*: 3.4 ±0.1 µm/h; p > 0.05; n = 719 cells from *Ccr2^−/−^Cx3cr1^gfp/+^* and 405 cells from *Ccr2^+/+^Cx3cr1^gfp/+^*mice; determined using an unpaired t-test). N = 4 *Ccr2^−/−^Cx3cr1^gfp/+^*and 4 *Ccr2^+/+^Cx3cr1^gfp/+^*mice. Note that the *Ccr2^+/+^* control conditions are the same measurements as the *Cx3cr1^gfp/+^* values in Figure 1.

## Discussion

So far, the influence of CX3CR1 on cellular migration has not been studied in detail. Quantitative global proteomic analysis suggested changes in migration-relevant signaling pathways of cells in the cortex. Hence, we performed *in vivo* two-photon imaging, to characterize the role of CX3CR1 in microglia migration, as one elementary feature of those immune cells. In prior studies, migration and relevant factors regulating migration were mostly studied *in vitro* and only a few studies focused on microglial migration *in vivo*. Specifically, the dynamics of the microglial response to brain injury remained elusive. Our *in vivo* study demonstrates that microglial migration induced by a laser lesion is modulated by CX3CR1. We found that the absence of CX3CR1 augments microglial velocity and distance travelled to the lesion. Similarly, extension of microglial processes towards the lesion is enhanced in the *Cx3cr1*-knockout mice. Our data show local proliferation of microglia accumulating around the lesion (∼2%). Lastly, by using *Ccr2*-knockout mice, we could exclude a contribution of peripheral monocyte-derived cells.

By analyzing the proteome of the cortex, we were able to identify a variety of pathways that are differentially regulated in the absence of CX3CR1. Although the proteomics results cannot provide information about individual cell types the results are likely to reflect microglial changes, because CX3CR1 primarily expressed on microglia in the cortex. Interestingly, amongst the identified DEPs many are related to cytoskeleton and migration of cells. This analysis reveals a novel functional link between CX3CR1 and the cytoskeleton, as well as cellular movement and migration. Most prominently, the IPA results from proteomic analysis imply a downregulation of RHO-signaling, the associated signaling processes regarding the cytoskeleton, and migratory capacity of the cells within the cortical tissue of the *Cx3cr1*- knockout mice. Initially, this result seemed counter-intuitive in light of our results of enhanced migration and fine processes extension. However, it has been shown that RhoA-GTPase deficiency in Cx3cr1^+^ cells resulting in ablation of Rho-signaling is associated with spontaneous microglia activation and increased expression of pro-inflammatory cytokines *in vivo*^22^. Additional *in vitro* results confirmed that a tight control of RhoA activity is necessary to control the microglial inflammatory response, since a complete Rho-KO led to microglial cell death, whereas a slight reduction led to increased inflammatory activity. In contrast, sustained RhoA signaling inhibited inflammatory microglial activity^25^. These results imply that a downregulation, but not total deactivation, of Rho-signaling induces microglial activation, which would make them more sensitive to an injury, such as a laser lesion. Thus, our finding of reduced RhoA-signaling could be associated with microglial activation, rendering them alert for a faster injury response. Another discovery is that the profile of altered DEPs predicted elevated motor dysfunction and movement disorders in the *Cx3cr1-KO* mice. Interestingly, our proteomics-based predictions concerning motor dysfunction have already been described comparing motor behavior of *Cx3cr1^+/–^*with *Cx3cr1^−/−^* mice^26,27^.

Microglial function relies on their motility and ability to migrate. By extending and retracting their processes as well as active migration towards sites of injury within the CNS, microglia exert a number of functions. We demonstrate that deficiency of the chemokine receptor CX3CR1 led to an increased migration velocity and travelled distance of microglia towards a laser lesion compared to *Cx3cr1*-haploinsufficiency. These findings are in accordance with previous research stating that under healthy conditions, a small amount of microglia is mobile. Focal vascular injury then induces migration towards the site of injury in a sex-specific manner^7,28^. We further found an increase in microglial density ∼140 μm around the lesion. Similar patterns of spatial microglial migration were found in the same mouse model upon microhemorrhage-induction^19^. In a similar manner, seizures as well as whisker trimming were found to induce microglia migration^28^. These studies utilized the same *Cx3cr1^gfp/+^* mice and clearly showed the onset of microglia migration upon different stimuli, however lack investigation of CX3CR1-dependency. Thus, we reveal a novel role of CX3CR1 in regulating characteristics of microglial migration *in vivo*. In a similar manner, but on a smaller timescale, processes extension and velocity towards a lesion are affected by CX3CR1-deficiency, which underlines the role of CX3CR1 in the fine-tuning of microglial motion in response to brain damage. However, under healthy conditions, our and a previous study found unchanged microglial motility comparing *Cx3cr1^+/–^* and *Cx3cr1^−/−^* knockout mice^17^, indicating that microglial processes are differentially regulated under unperturbed and injury conditions. In line with this, highly directional process extension of microglia towards ATP release, as a result of brain tissue damage, has been described *in vivo*^8^. The link between ATP and microglia movement was further underlined in a study investigating the effect of the P2Y12-receptor knockout, which found that in these mice microglial rearrangement is significantly reduced^28^. In our model, *Ccr2*- deficiency did not affect microglia migration. In addition, similar numbers of accumulating cells at the lesion in *Ccr2-*deficient mice exclude a contribution of CX3CR1^+^CCR2^+^ monocytes from the blood. The *Ccr2*-independency of microglial migration also implies that its ligand MCP-1 does not play a major role in regulating microglia migration in the context of brain lesions.

Occasionally, we have observed newly appearing cells around the lesion due to local proliferation. Proliferation of microglia has been proposed and described over longer observation periods, in which fluorescently-labelled microglia were imaged over the course of many weeks. Putative division events around amyloid lesions, similarly to what we found, was sometimes preceded by an increase in soma-volume^29^. However, the fusion or simply neighboring of two microglia might easily misinterpreted as a division event. Therefore, definite proof of divisions observed with imaging methods requires either higher spatio-temporal resolution or the combination of imaging with DNA-intercalating markers like EdU and *post-mortem* retrieval of the same cells using immunohistochemistry. Here, we were able to prove local microglia proliferation after migration towards a laser lesion using EdU as a proliferation marker. Microglia proliferation around an infarct in the brain parenchyma was also shown in prior studies, using EdU as a proliferation marker^19^. Therefore, proliferation on a much smaller level than migration, also contributes to local accumulation around a laser lesion.

Based on our findings, we propose a model for microglial kinetics upon brain injury: Within the first hour after a laser lesion, microglial processes extend towards the lesion, which is followed by a highly directed migration towards the lesion. Migration, peaks at approximately 16-24h post lesion-induction and, lastly, approximately 48h after lesion-induction, a small fraction of the microglia that migrated towards the lesion start to proliferate there (**Fig. 6**). Our findings are in line with previous results, that propose a similar cascade of events during microglia migration^30^.

**Figure 6.**
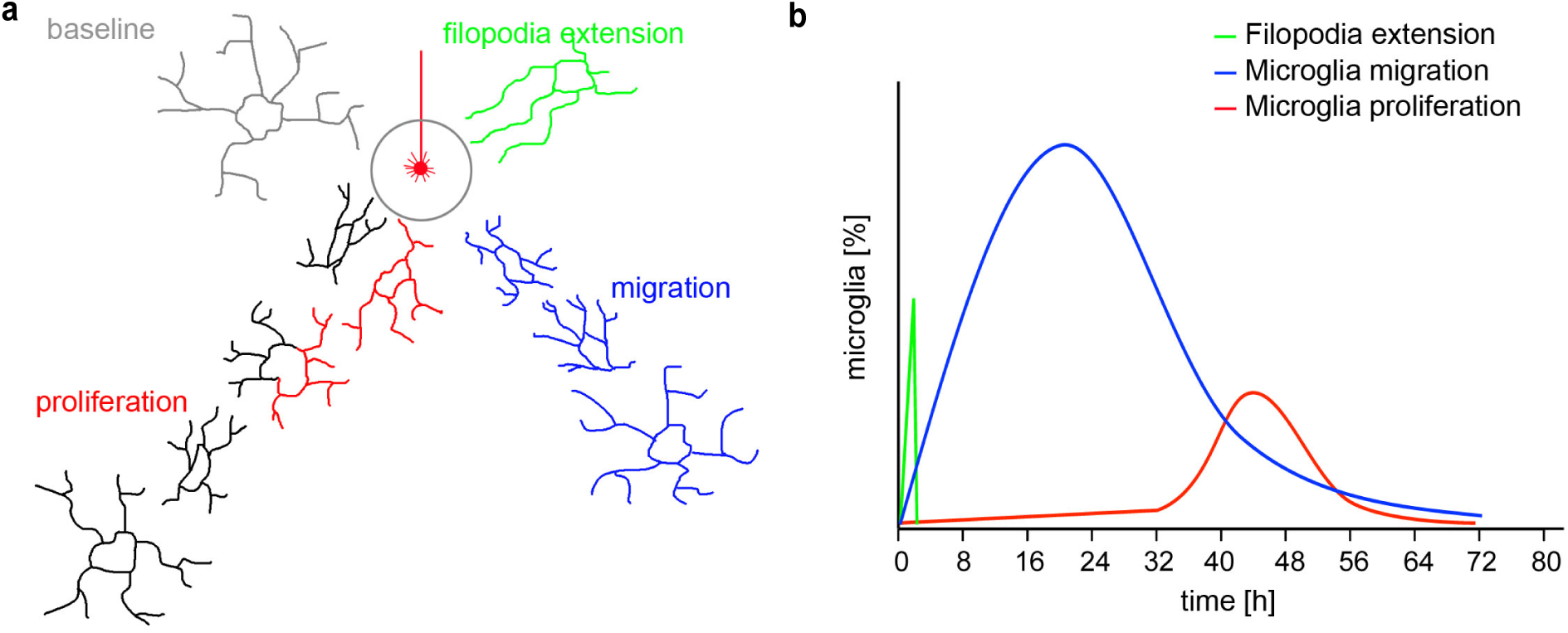
Schematic of microglia expansion kinetics. **(a)** Under baseline conditions (grey), microglia remain at their position and are constantly screening the brain parenchyma by protracting and retracting their fine processes. Upon brain injury, microglia fine processes extend within minutes towards the lesion (green). Subsequently, about 25% of the resident microglia in the brain parenchyma migrate towards the lesion (blue). In addition to migration, an even lower fraction of about 2% of microglia migrates and proliferates in proximity to the lesion (black and red). **(b)** Proposed model of kinetics and amount of fine processes extension (green), microglia migration (blue) and proliferation (red) starting at time-point 0 (lesion-induction) till 72h-post lesion.

Whether microglial activation upon brain injury is beneficial for tissue repair or if it hinders healing is still not fully elucidated. It is known that microglia are key neuroprotective factors after spinal cord injury forming a glial scar^31^ and also exert neuroprotective functions as their depletion leads to a severely increased infarct size^32–34^. Contrastingly however, growing glial scars were also found to impair axonal regeneration^35^ and microglia release glutamate, which is a major excitotoxic factor^36^. Hence, microglia seem to have a dichotomous role in response to CNS injury. We found that microglia, in mice of all different genotypes used, are recruited towards laser lesion-induced damage and therefore are direct responders to injury. Microglia migration towards the site of injury is thus the first step of a complex process of the microglial injury response and precedes their effector functions. Therefore, modulating migration via CX3CR1 might represent a novel strategy for the treatment of diseases including a sterile innate immune response.

## Methods

### Transgenic mice

For label free proteomic analysis, *Cx3cr1^CreERT2^* mice that featured a homo- (*Cx3cr1^−/−^*) or heterozygous (*Cx3cr1^+/–^*) knockout of the fractalkine receptor, replaced with a CreERT2-Recombinase were used. Animals were 14.6 ±0.3 months old and sexes were equally distributed within groups. For the in vivo imaging experiments, *Cx3cr1^gfp^* mice at 5.4 ±0.7 months and with equally distributed sexes were used. Both mouse lines were generously provided by Steffen Jung (Weizmann Institute of Science, Rehovot, Israel)^37^. Homozygous *Cx3cr1^gfp/gfp^* mice were crossed with heterozygous *Cx3cr1^gfp/+^* mice to yield homozygous and heterozygous littermates at a 1:1 ratio. The heterozygous *Cx3cr1^gfp/+^*mice retain a significant portion of the CX3CR1 receptor, while *Cx3cr1^gfp/gfp^ mice* feature a complete *Cx3cr1* knockout. Genotyping of the mice was performed using the following primers: IMR3945 5’-TTCACG TTCGGTCTGGTGGG-3’, IMR3946 5’-GGTTCCTAGTGGAGCTAGGG-3’, IMR3947 5’-GAT CACTCTCGGCATGGACG-3’. *Cx3cr1^+/+^* yield a single ∼970 bp fragment, *Cx3cr1^gfp/+^* yield a ∼1200 bp and a ∼970 bp fragment, and *Cx3cr1^gfp/gfp^* a single ∼1200 bp fragment. *Cx3cr1^gfp/+^Ccr2^−/−^* mice were generated by Daniel R. Engel and Christian Kurts by crossing *Cx3cr1^+/+^Ccr2^−/−^* with *Cx3cr1^gfp/gfp^Ccr2^+/+^*mice yielding heterozygous *Cx3cr1^gfp/+^Ccr2^+/–^* offspring that was back-crossed with *Cx3cr1^+/+^Ccr2^−/−^* mice^38^. Since both, *Cx3cr1* and *Ccr2,* are located on the murine chromosome 9 we had to identify a crossing over event. *Cx3cr1^gfp/+^Ccr2^−/−^*mice were selected by levels of *Ccr2* and *Gfp* detected via Flow Cytometry of tail vein blood and genotyping using the following primer: P1: 5’-TAA ACC TGG TCA CCA CAT GC-3’; P2: 5’-TCA CTT ACT TTA CAA CCC AAC C-3’; P3: 5’-GGA GTA GAG TGG AGG CAG GA-3’; P4: 5’-GAA GGC GAT AGA AGG CGA T-3’. P1 and P2 amplify a 1.4 kb fragment of the wild-type allele and a 3.2 kb fragment of the knock-in allele, while P2 is located in a region that was not part of the original target allele, thereby checking for correct insertion. P1 and P3 amplify a 494 bp fragment of the wild-type allele. P3 spans the BamH1 restriction site used for neomycin gene insertion. P1 and P4 amplify a 1.2 kb fragment and P4 is located inside the neomycin gene (**Supplementary** Fig. 4).

Mice were group-housed in individually-ventilated cages (IVC) under specific pathogen free (SPF) conditions with *ad libitum* access to food and water. Mice of both sexes were used. After surgery, the mice were separated and singly housed. All procedures were in accordance with an animal protocol approved by the DZNE and the government of North-Rhine-Westphalia (Az 84-02.04.2017.A098).

### Label-free quantitative proteomics analysis

1 mg of mouse brain cortex from each genotype was dissected and processed using a modified filter-aided sample preparation method as described^39^. The peptide samples were cleaned using C18-reverse-phase ZipTipTM (Millipore). Dried peptide digest was resuspended in 1% trifluoroacetic acid and sonicated in a water bath for 1 minute before injection. Fractionated protein digests were analyzed in nano-LC–Thermo Q Exactive Plus Orbi-Trap MS with parameters set as described^40^.

Following LC-MS/MS acquisition, the raw files were qualitatively analyzed by Proteome Discoverer, version 2.5 (Thermo Scientific). The identification of proteins by Proteome Discoverer was performed against the UniProt/SwissProt reviewed mouse protein database (release 10-2022 with 17139 entries) using the built-in SEQUEST HT engine. The following parameters were used: 10 ppm and 0.02 Da tolerance for MS and MS/MS, respectively. Trypsin was used as the digesting enzyme, and two missed cleavages were allowed. The carbamidomethylation of cysteines was set as a fixed modification, whereas the oxidation of methionine and deamidation of asparagine and glutamine were set as variable modifications. The false discovery rate was set to less than 0.05, and a minimum length of six amino acids (one peptide per protein) was required for each peptide hit. The normalized LFQ intensities were analyzed with Perseus software, version 2.0.11 (https://maxquant.net/perseus/). The MQ data were filtered to exclude biological and statistical contaminants. Only proteins detected in 70% in each sample group were taken into account in the quantitative analyses. After data filtering and transformation to log2(x), the missing values from normal distribution were imputed. The fold changes in the level of proteins were assessed by comparing the mean intensities among both experimental groups. A protein was considered to be differentially expressed if the difference between 2 groups was statistically significant (unpaired t-test with permutation-based FDR <0.05), and the fold change of 1.5. Only DEPs identified with a minimum of 2 unique peptides, at >99% confidence level were accepted. Pathway analysis was done using Ingenuity Pathway Analysis™ (QIAGEN), to determine canonical pathways, and disease and function annotations associated with the observed differences in protein profiles. The prediction of the direction of change for a given function or pathway was estimated by a z-score ≥2 (for activation) and ≤-2 (for inhibition)^41,42^.

### Cranial window surgery for cortical window implantation

We surgically implanted an open skull cranial window over the right cortical hemisphere as previously described (Bittner et al., 2010; Druart et al., 2021; Fuhrmann et al., 2007, 2010; Holtmaat et al., 2009). Mice were anesthetized via intraperitoneal (i.p.) injection of ketamine/xylazine (0.13/0.01 mg/g body weight). Additionally, dexamethasone (0.2 mg/kg) and TEMGESIC^®^ (0.05 mg/kg; Reckitt Benckiser Healthcare (UK) Ltd., Great Britain) were administered i.p. prior to surgery. Under anesthesia, we removed a circular piece of the skull over the somatosensory cortex (5 mm diameter) using a dental drill (Schick-Technikmaster C1; Pluradent; Offenbach, Germany). Special care was taken to leave the dura intact. The dura was covered with PBS immediately to prevent it from drying out. After the craniotomy, we glued a circular coverslip (5 mm diameter, thickness #01) to the skull using dental acrylic (Cyano-Veneer fast; Heinrich Schein Dental Depot, Munich, Germany) to close the open area. A small metal head-bar with a screw thread was glued next to the coverslip to allow repositioning of the mouse during longitudinal imaging sessions. The imaging experiments were started after a three-week resting period following cranial window surgery to ensure a stable window and to prevent neuroinflammation.

### Two-photon *in vivo* imaging

Microglia migration imaging was performed using an upright Zeiss (Carl Zeiss Microscopy GmbH, Jena, Germany) Axio Examiner LSM7MP setup, equipped with a Coherent Cameleon Ultra2 two-photon laser (Coherent, Dieburg, Germany). Microglia fine processes extension analysis was carried out using a TriM Scope II (LaVision BioTec GmbH, Bielefeld, Germany) equipped with a Coherent Cameleon Ultra2 two-photon laser. Green fluorescent protein (GFP) was excited at 920 nm, the fluorescence emission was bandpass filtered (525/50) and detected by an ultra-sensitive non-de-scanned gallium-arsenide-phosphide (GaAsP) detector with large aperture detection optics in close proximity to the objective. A Zeiss 20x water immersion objective (NA1.0) or a Nikon 16x water immersion objective (NA0.8) were used for magnification. The standard microscopy table was replaced by a custom-made table to allow repeated positioning of a living mouse under the microscope. At each imaging time-point, mice were briefly anesthetized with 0.8% Isoflurane gas anesthesia that lasted for 15-20 minutes. Image acquisition was performed with the Zeiss ZEN2010 software or Imspector, respectively. To follow the migration or fine processes extension of individual microglia over time, we acquired z-stacks, 300 µm in size, starting from the cortical surface with a 3 µm z-resolution. Image frames were 600 x 600 µm in size with 1536 x 1536 pixels in x,y-direction (migration) or 1243 x 1243 pixels (fine processes extension). At the first imaging time-point, the distribution of microglia was determined by acquiring a z-stack as described. Subsequently, a laser lesion was induced by focusing the laser beam on the middle position in x, y- and z-dimension (x: 300 µm, y: 300 µm, z: 150 µm). The wavelength was changed to 800 nm and an output power of 180 ±5 mW. An area of 10 µm x 10 µm was scanned for 1.5 sec with a pixel dwell time of 1.27 µsec and 0.01 µm/pixel pixel size. Subsequently, z-stacks were acquired every 5 minutes for fine processes extension analysis and at 4-6 hours over a period of 72 hours for cell body migration analysis. Care was taken to ensure similar fluorescence levels in space and time. Repositioning over time was achieved by orienting to the vascular pattern under reflected light illumination using a GFP filter set and a metal halide lamp HXP100 (Carl Zeiss Microscopy GmbH, Jena, Germany). Additionally, orienting at the center of the lesion in x-, y- and z-dimension was carried out using two-photon illumination.

### Retrieval of proliferating microglia in fixed tissue after *in vivo* imaging

To retrieve previously imaged migrating and proliferating microglia after *in vivo* imaging in fixed tissue, mice were injected with EdU (5-ethynyl-2’-deoxyuridine; 25 µg) starting at the first imaging time-point every 12 hours until the end of imaging after 72 hours. Immediately after the 72-hour imaging time-point, mice were transcardially perfused with phosphate buffered saline (PBS) followed by 4% paraformaldehyde (PFA). The brain was left inside the intact scull and only the circular coverslip was removed to give access of PFA to the brain for post-fixation over night at 4°C. On the next day, the mouse head was fixed via the metal head-bar (see cranial window surgery) in the same plane as imaging had been carried out. The scull over the right cortical hemisphere was carefully removed by using a dental drill. Subsequently, the mouse head was positioned under a vibratome blade (Leica VT1000S, Wetzlar, Germany) to cut the brain in 100 µm-thick sections. The sections were oriented in the same plane as the imaging had been carried out to generate sections perpendicular to the z-focusing axis of the microscope. Four to five sections were cut and successively collected in individual wells of a 24-well plate. Care was taken to keep the topside of the slices up in the PBS-filled well.

Immunohistochemistry was carried out with a slightly adapted protocol from Gogolla and colleagues^46^: The sections were permeabilized with 2% Triton X-100 for 12 hours at 4°C on a shaker. Slices were washed with 3% BSA in PBS three times for 5 minutes per wash. The Click-IT^®^ reaction cocktail was prepared as described in the manufacturer’s manual (Click-IT^®^ EdU Alexa-Fluor^®^ 647 Imaging Kit, Invitrogen) and 500 µl of the solution were added to each well. Slices were incubated for a period of 60 minutes protected from light. Subsequently, the reaction solution was removed and the slices were washed with 3% BSA in PBS. This procedure avoided an additional staining of microglial *Gfp*, since the very mild staining conditions keep *Gfp* intact. After the staining, slices were mounted on glass objective slides using Fluoromount as mounting medium and glass coverslips. Brain slices were analyzed under epi-fluorescence using a GFP filter set. Slices that contained lesions were identified and confocal laser scan images were acquired from the lesion on an inverted Zeiss LSM700 microscope equipped with a 488 nm Ar-ion and 635 nm HeNe laser to excite GFP and Alexa647. Fluorescence was detected at 500-550 nm (GFP) and at >650 nm (Alexa 647). A Zeiss 20x objective with NA0.8 or 40x immersion oil objective NA1.3 was used to acquire z-stacks of microglia around the lesion (Carl Zeiss Microscopy GmbH, Jena, Germany). Images with a frame size of 2048 x 2048 pixel, a pixel size of 0.2 µm/pixel and a z-step of 1 µm were acquired in two channels (GFP and Alexa-Fluor 647). Re-localization of proliferating microglia was carried out by manually scrolling through the 72-hour *in vivo* imaging stack and the confocal image stack covering the whole lesion in x, y and z-dimension in parallel.

### Image processing and data analysis

For fine processes extension analysis, images were processed with the open-source software Fiji^47^. Ome.tiff files from ImSpector (LaVision Biotec, Bielefeld, Germany) were median filtered and maximum intensity projected. Images were aligned in space and time using the “StackReg” plugin in Fiji^48^. Individual fine proecesses were tracked using the Fiji plugin “Manual Tracking” by Fabrice Cordelières, Institut Curie, Orsay (France). The lesion diameter, the Euclidean distance from start to end position and the total distance that the fine processes traveled were measured 45 minutes post lesion. The fine processes velocity was calculated as distance traveled divided by time. The meandering index (MI) was calculated by dividing the Euclidean distance by the distance traveled 45 minutes post lesion.

Microglia cell body migration was analyzed using ZEN, Fiji and Volocity as analysis software. After acquisition with ZEN, images were median filtered. Subsequently, images were aligned in z,t-dimension by manually aligning the top and bottom image for each time-point. Twenty slices, spanning 60 µm and containing the lesion, were maximum intensity projected using the depth color-coding function in ZEN. Microglia at the bottom were colored in blue and microglia at the top of the projection were colored in red (**Supplementary** Fig. 2). These images were exported to Fiji and aligned in x,y-dimension over the whole imaging time-period using the “StackReg” plugin. The resulting time-series were exported to Volocity (Volocity Ver. 5.5, Perkin Elmer, Rodgau, Germany), which was used for tracking individual microglia. Microglia identified as dividing during tracking were re-localized in fixed brain sections and tested for EdU-positivity (see “Retrieval of proliferating microglia in fixed tissue after *in vivo* imaging”). Since imaging was carried out every 4-6 hours, all time-dependent data was binned in eight-hour bins. The fraction of moving cells was calculated by dividing the number of moving cells at a given time-point by the total number of cells in the field of view of the 60 µm maximum-intensity projection. To calculate the distance traveled of microglia towards the lesion, the average distance traveled of all moving cells (translocation > 4 µm) at a given time-point was calculated. The velocity was calculated by dividing the distance traveled [µm] of all migrating microglia (translocation > 4 µm) by 72 hours. The fraction of proliferating microglia was calculated by dividing the number of dividing cells by the total number of cells in the field of view of the 60 µm maximum-intensity projections. The distance to lesion when microglia divided was measured as the Euclidean distance to the center point of the lesion. The MI of migrating microglia was calculated as the Euclidean distance divided by the distance traveled with the lesion center as reference point.

Microglia motility analysis was performed as previously described^17,49^. Image stacks were registered using the StackReg-Plugin in ImageJ^48^ and a maximum intensity projection was generated. The projection of all cells within this stack and their branches was merged with the projections from all recorded timepoints (15 timepoints per recording, delta t = 5 min) and registered with the Stack-Reg Plugin again. Subsequently, the time-points were pseudo-colored in red and green and the turnover rate of microglial processes was calculated for every time-point (**Supplementary** Fig. 3).

Data analysis was performed in a blind manner by an experimenter without knowledge of the experimental conditions. Statistical significance was assumed at *p* < 0.05. Statistical differences in measurements over time were determined using two-way ANOVA in the software GraphPad Prism, version 10.2.2 (GraphPad Software Inc., La Jolla, USA). Statistical differences between multiple different experimental groups were carried out by Bonferroni post-tests. For two-group comparisons, Student’s t-test was used.

## Data Availability Statement

All proteomics data generated or analyzed during this study are included in this published article and its Supplementary Information Files. Mass spectrometry data were deposited in the Mass Spectrometry Interactive Virtual Environment (MassIVE) database under accession number MSV000095652. Microscopy data will be deposited and openly accessible at DRYAD.

### Acknowledgements

This work was supported by the DZNE, and grants to MF by the European Union ERC-CoG (MicroSynCom 865618), ERA-NET Neuron (MicroSchiz) and the German research foundation DFG (SFB1089 C01, B06; SPP2395). The iBehave network funded by the Ministry of Culture and Science of the State of North Rhine-Westphalia to MF. KL, RS, and ML were supported by the Academy of Finland grant #318857. This research was also funded by the ERA-NET Co-Fund project JTC2017, MicroSynDep Consortium to KL, RS, ML and MF Proteomics measurements were done at the Meilahti Clinical Proteomics Core Facility (supported by HiLIFE and Biocenter Finland). HA is supported by a Boehringer Ingelheim Fonds PhD Fellowship.

## Author Contributions

JW, CH, HA and MF prepared figures and wrote the manuscript with input from all authors. JW, CH, MF, LS and KK carried out surgeries and recorded in vivo imaging data. SC and JS performed colony management and generated tissue samples. JS, JW and CH performed histology and microscopic analysis. RS, KL and ML acquired proteomics data. JW, CH, HA and ML analyzed data. CK, DE and JH provided lab-space, reagents and discussed the manuscript. ML and MF conceived, planned and supervised the project.

