## Supplemental Information for "CX3CR1 modulates migration of resident microglia towards brain injury"

### Supplementary Material

#### Supplementary Figures

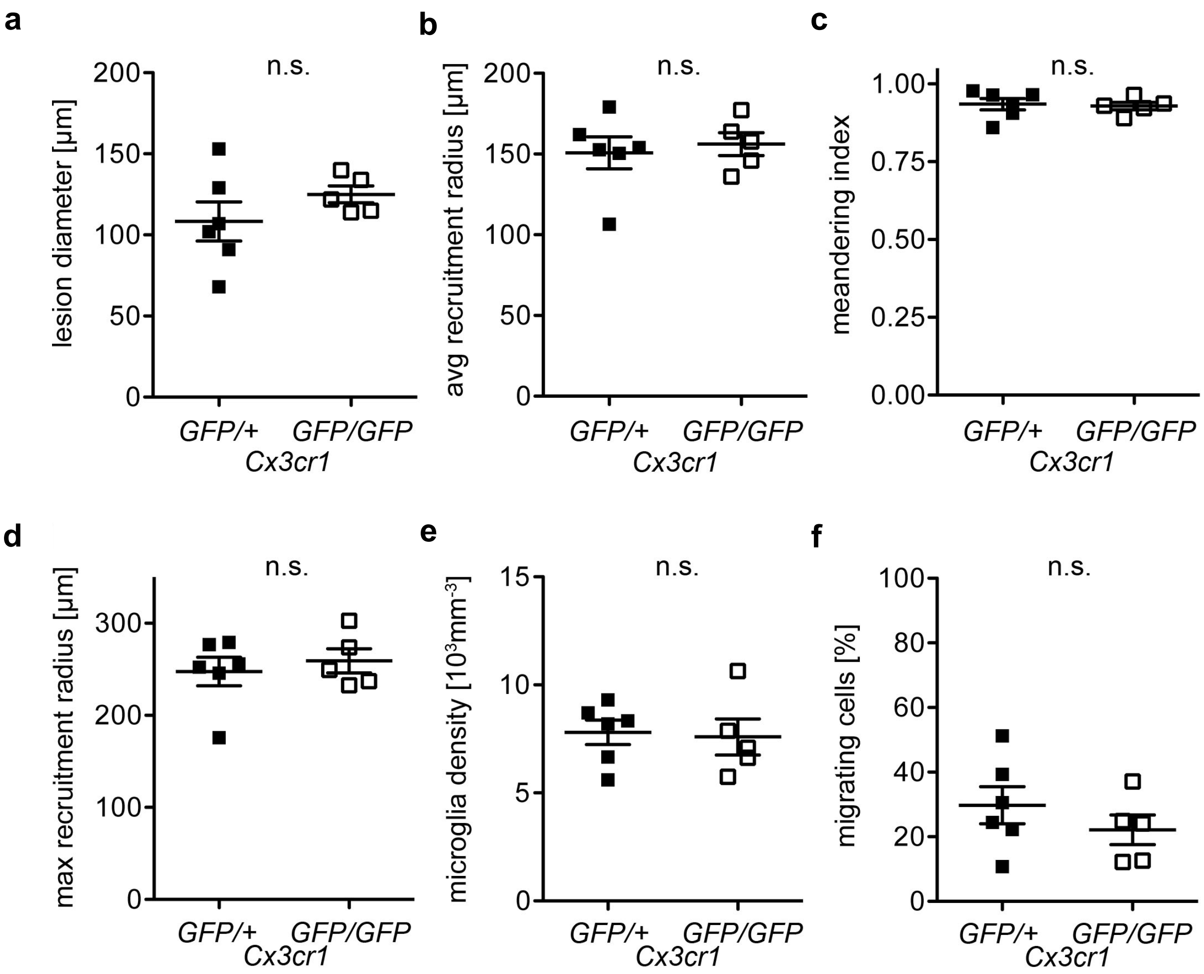


**Supplementary Figure 1. CX3CR1-independent parameters.**
**(a)** Average lesion diameter of *Cx3cr1*$^{gfp/+}$ and *Cx3cr1*$^{gfp/gfp}$ mice is unchanged (*gfp/*+: 108.3 ±12.1 µm; *gfp/gfp*: 125.0 ±5.2 µm; n = 5-6 mice/group, p > 0.05). **(b)** Unaltered average recruitment radius of migrating microglia comparing *Cx3cr1*$^{gfp/+}$ and *Cx3cr1*$^{gfp/gfp}$ mice (*Cx3cr1*$^{gfp/+}$: 150.8 ±9.8 µm; *Cx3cr1*$^{gfp/gfp}$: 156.2 ±7.1 µm; n = 5-6 mice/group, p > 0.05). **(c)** The meandering index of microglia migrating to the lesion reveals a high degree of directionality and was unchanged between *Cx3cr1*$^{gfp/+}$ and *Cx3cr1*$^{gfp/gfp}$ mice (*Cx3cr1*$^{gfp/+}$: 0.935 ±0.018; *Cx3cr1*$^{gfp/gfp}$: 0.929 ±0.012; n = 5-6 mice/group, p > 0.05). **(d)** Unchanged maximum recruitment radius between *Cx3cr1*$^{gfp/+}$ and *Cx3cr1*$^{gfp/gfp}$ mice (*Cx3cr1*$^{gfp/+}$: 248 ±15 µm; *Cx3cr1*$^{gfp/gfp}$: 259 ±13 µm; n = 5-6 mice/group, p > 0.05). **(e)** Microglia density before the laser lesion was unchanged between *Cx3cr1*$^{gfp/+}$ and *Cx3cr1*$^{gfp/gfp}$ mice (*Cx3cr1*$^{gfp/+}$: 7801 ±567 µm; *Cx3cr1*$^{gfp/gfp}$: 7593 ±838 µm; n = 5-6 mice/group, p > 0.05). **(f)** Fraction of migrating cells in the analyzed volume is not significantly different between the two genotypes (*Cx3cr1*$^{gfp/+}$: 29.7 ±5.7%; *Cx3cr1*$^{gfp/gfp}$: 22.2 ±4.6 %; n = 5-6 mice/group, p > 0.05).

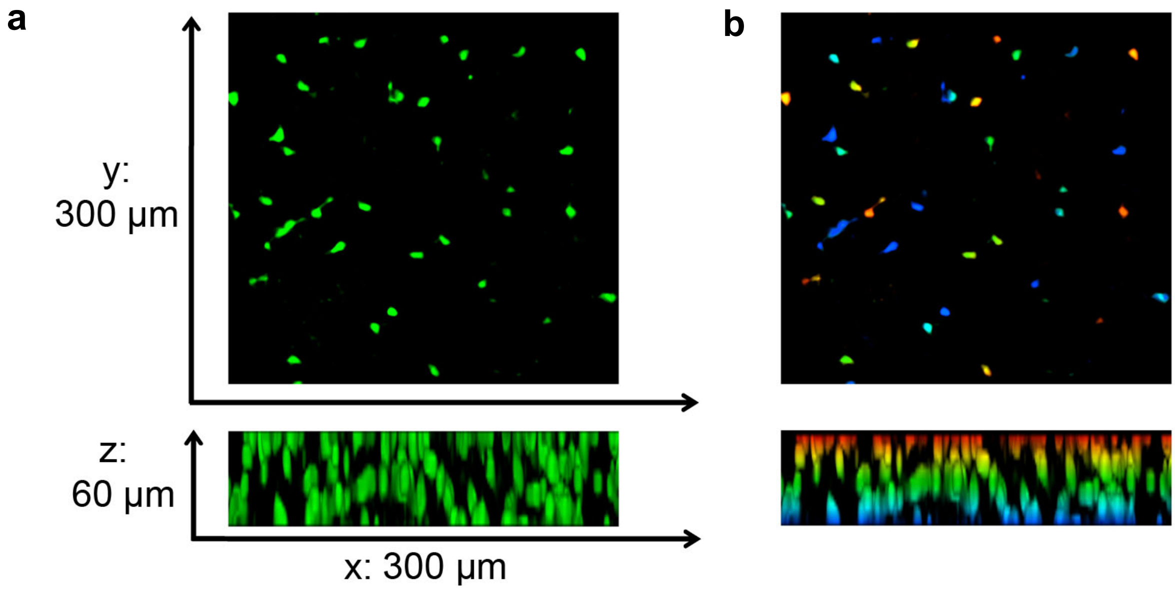


**Supplementary Figure 2. Color-coded depth projection.**
**(a)** Maximum intensity projection of a 60 µm depth-spanning z-stack containing GFP-labeled microglia acquired with a z-step of 3 µm distance. Upper panel x, y-projection lower panel x, z-projection. **(b)** Color-coded maximum intensity projections (x, y and x, z) of the same z-stack. Different depths of individual microglia cells are clearly visualized.

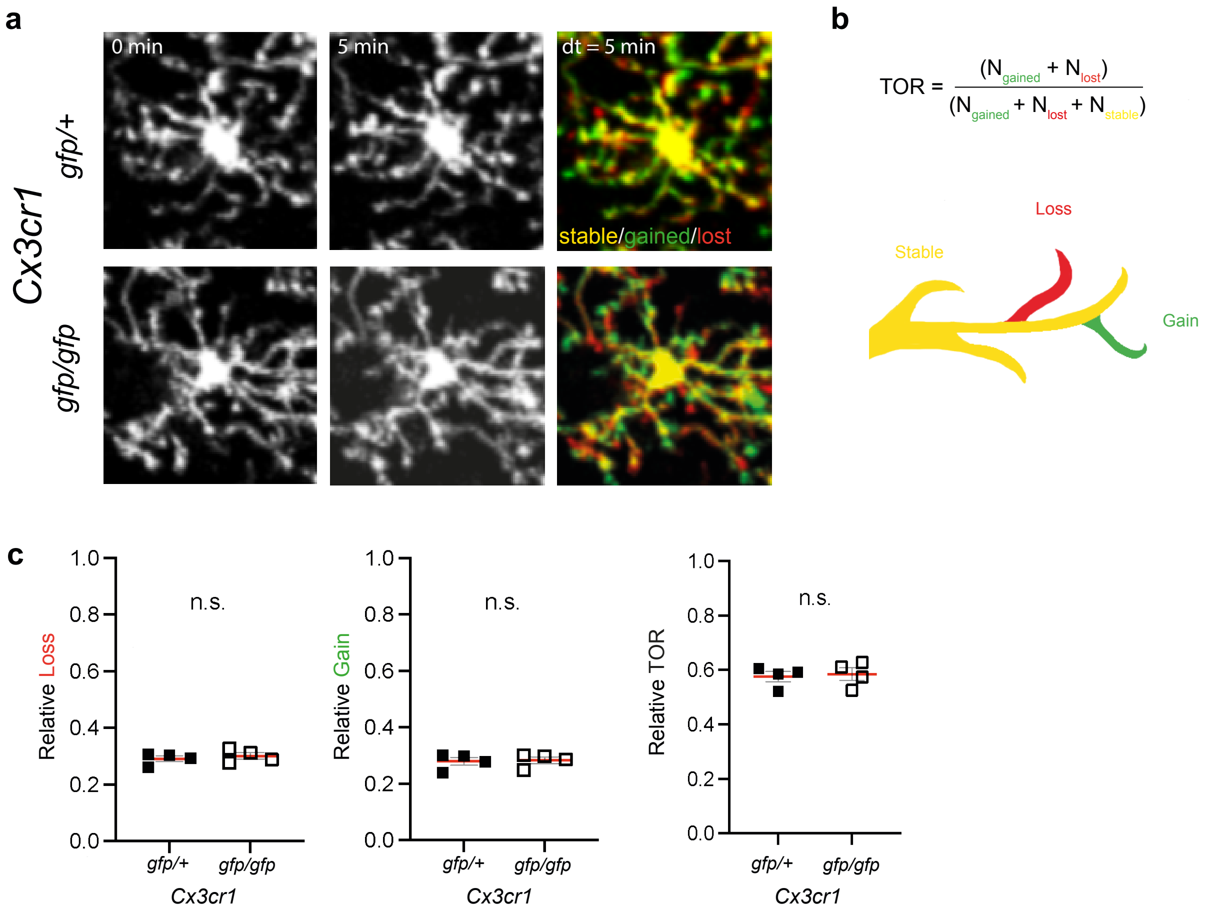


**Supplementary Figure 3. Microglia Fine Processes Turnover Rate**
**(a)** Exemplary images of microglia fine processes turnover in the cortex comparing *Cx3cr1*$^{gfp/+}$ and *Cx3cr1*$^{gfp/gfp}$ mice. **(b)** Microglia fine processes turnover rate (TOR) was calculated as the number (absolute pixel value) of lost, $N_{lost}$ (red), and newly gained, $N_{gained}$ (green) pixels divided by the sum of all pixels within a region of interest. **(c)** Microglia fine processes gain, loss and turnover rate of *Cx3cr1*$^{gfp/+}$ and *Cx3cr1*$^{gfp/gfp}$ mice (n = 4 stacks per mouse, N = 4 mice/group), unpaired t-test, two-tailed). Error bars: SEM.

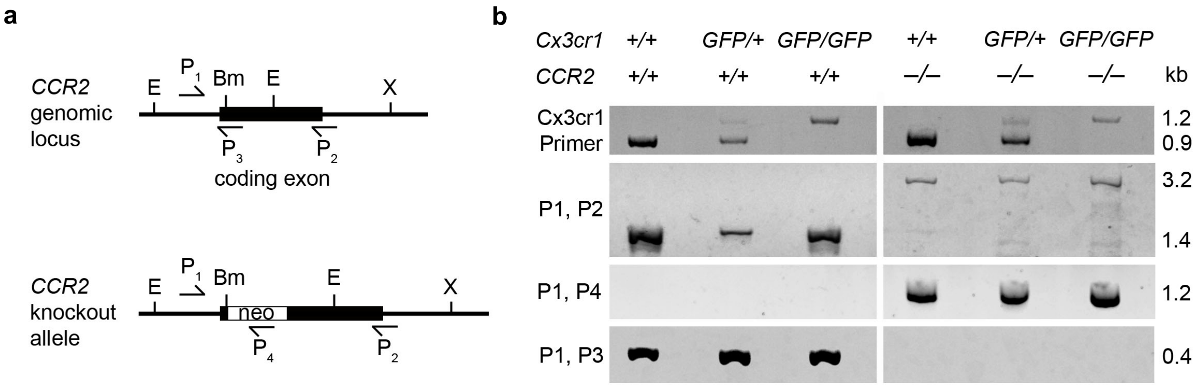


**Supplementary Figure 4. Generation of *Ccr2*$^{-/-}$*Cx3cr1*$^{gfp/gfp}$ mice.**
**(a)** Illustration of the genomic *Ccr2* locus and the *Ccr2* knockout allele with various restriction sites EcoR1 (E), BamH1 (Bm), Xba1 (X) and Primer location P1-P4. Primer P1 lies in the non-coding region before the ATG start-codon. P2 is located at the exon-intron boundary. This part was not included in the targeting construct [18]. P3 is located at the BamH1 restriction site, where the neomycin resistance was introduced. P4 is part of the neomycin resistance. **(b)** Exemplary images of agarose gels for the different genotypes of *Ccr2::Cx3cr1*$^{gfp}$ mice. To determine the zygosity for *Cx3cr1*$^{gfp}$, the PCR resulted in a 970 bp fragment for the wild-type allele and a 1.2 kb fragment for the knockout allele. Heterozygous mice display both bands. To determine *Ccr2* zygosity, primer P1-4 were used. P1 and P3 amplify a 400 bp exclusively from the wild-type allele, while P1 and P4 yield a 1.2 kb fragment exclusively from the *Ccr2* knockout allele. Primer P1/P2 amplify a 1.4 kb and a 3.2 kb fragment from wild-type and knockout allele respectively. Moreover, P2 is located at the exon-intron boundary of the *Ccr2* gene that was not part of the targeting construct.

Supplementary Material

#### Supplementary Videos

**Supplementary Video 1. Microglia Migration.**
Time-lapse video of microglia migration after a laser lesion over a period of 72 hours. In the left panel, microglia are depicted in green. The right panel shows individual microglia cells as white dots moving towards the lesion over the entire imaging period. Interestingly, previously sessile microglia migrate towards the lesion. Scale bar: 50 µm.

**Supplementary Video 2. Microglia Filopodia Extension.**
Microglia fine-process extension upon induction a laser lesion within the first 75 minutes acquired in five-minute time-intervals. The left panel shows an overview image of microglia in close proximity and further away from the lesion spot. The right panel shows a zoom-in of the time-lapse of individual microglia fine processes extension. The blue line exemplarily illustrates a tracked microglial fine process. Scale bars: 35 µm (left), 10 µm (right).

**Supplementary Video 3. Microglia Division.**
Exemplary *in vivo* time-lapse of a dividing microglial cell after a laser lesion over a time period of 72h. The brain slices were fixed in PFA after 72h. *In vivo* imaged microglia (top right) were re-localized in the fixed brain slices and immunohistochemical staining was performed for the proliferation marker EdU (red, bottom left). Additionally, GFP in microglia is displayed (green, top right) and the images are depicted individually and merged (bottom right) in order to show EdU-positive proliferating cells. Red and green arrows indicate dividing microglia, the blue arrow indicates a non-dividing microglial cell. Scale bar: 25 µm.

#### Supplementary Tables

**Supplementary Table 1. All identified differentially regulated proteins in the cortex of *Cx3cr1*$^{-/-}$ and *Cx3cr1*$^{-/+}$ mice.**
**(1)** All 2668 identified proteins are depicted including their accession numbers, description of the protein function, False Discovery Rate (FDR), q-values, Posterior Error Probability (PEP) Score, Coverage, Peptide number out which the number of unique peptides for each protein was identified and information of the protein characteristics. **(2)** From all detected proteins, 168 proteins that are differentially expressed between both conditions were identified. Apart from the protein characteristics, the table displays the statistical analysis conducted via the Student's t-test.

**Supplementary Table 2. Ingenuity Canonical Pathways Analysis.**
Based on the previously identified differentially regulated proteins, Ingenuity Pathway Analysis (IPA) was conducted to predict pathways that are altered in the cortex of the *Cx3cr1*$^{-/-}$ compared to the *Cx3cr1*$^{-/+}$ mice.
